# The von Willebrand Factor stamps Plasmatic Extracellular Vesicles from Glioblastoma Patients

**DOI:** 10.1101/2021.03.25.437073

**Authors:** Quentin Sabbagh, Gwennan André-Grégoire, Carolina Alves-Nicolau, Nicolas Bidère, Emmanuel Jouglar, Laëtitia Guével, Jean-Sébastien Frénel, Julie Gavard

## Abstract

Glioblastoma is a devastating tumor of the central nervous system characterized by a poor survival and an extremely dark prognosis, making its diagnosis, treatment and monitoring highly challenging. Numerous studies have highlighted extracellular vesicles (EVs) as key players of tumor growth, invasiveness and resistance, as they carry and disseminate oncogenic material in the local tumor microenvironment and at distance. However, whether their quality and quantity reflect individual health status and changes in homeostasis is still not fully elucidated. Here, we separated EVs from plasma collected at different time points alongside with the clinical management of GBM patients. Our findings confirm that plasmatic EVs could be separated and characterized with standardized protocols, thereby ensuring the reliability of measuring vesiclemia, *i.e*. extracellular vesicle concentration in plasma. This unveils that vesiclemia is a dynamic parameter, which could be reflecting tumor burden and/or response to treatments. Further label-free liquid chromatography tandem mass spectrometry unmasks the von Willebrand Factor (VWF) as a selective protein hallmark for GBM-patient EVs. Our data thus support the notion that EVs from GBM patients showed differential protein cargos that can be further surveyed in circulating EVs, together with vesiclemia.

## Introduction

Glioblastoma is the most common primary malignant brain tumors in adults, and the most aggressive one, making it a major therapeutic challenge. Occurring in 70% of the cases between 45 and 70 years-old patients, prognosis remains extremely poor despite standardized, combative treatment. Indeed, tumor relapse is almost inevitable 7 to 10 months post-therapy, while the median survival is estimated at 14 months and the 5-year survival rate is about 5%. The actual frontline standard reference treatment has been established by Stupp *et al*. in 2005 and combines debulking surgery (if possible), followed by a six-week adjuvant radiochemotherapy followed with a six-month chemotherapy, both based on standardized doses of temozolomide (TMZ), an alkylating agent (1). Despite this harsh therapeutic regimen, glioblastoma recurrence is rapid and fatal. In this context of therapeutic impasse, new therapeutic strategies are needed.

Numerous studies have unveiled the central role of Extracellular Vesicles (EVs) as key mediators of intercellular communication in the glioblastoma microenvironment (2–8). The term EVs is a moniker for a variety of small, heterogeneous, membrane vesicles (30-1000nm), released by virtually all cells into surrounding milieu and consecutively circulating in biofluids, such as blood, urine *etc*… (9). These lipid bilayer vesicles transport protected bioactive materials *i.e*. proteins, lipids, and nucleic acids from their cell of origin to recipient cells, located either in the vicinity or at distance (10). EVs therefore act as potent mediators for cell communication in tissue and throughout fluids and body. Indeed, the EV secretory pathway is suspected to be perverted by both tumor and stromal cells, and might distribute oncogenic material and non-physiological information (2, 5, 6, 8, 10, 11). Thereby, EVs have been suggested to serve as major communication tools from and towards stem-like tumor cells, differentiated tumor cells, immune system and vascular endothelial cells, to support tumor growth, invasiveness and survival (2). Beside their instrumental role in transcellular message delivery, EVs quantitatively and qualitatively may vary with gender, ageing and during diseases (12, 13). EVs may thus represent a valuable, informative resource on health status and disease progression. Vesiclemia, *i.e*. EV quantity in a given volume, can be obtained from minimally invasive liquid biopsies and is currently scrutinized in a wide variety of diseases (12). However, whether vesiclemia reflects health status and changes in homeostasis is still not fully elucidated. Here, we established reproducible, standardized methodologies to isolate EVs from plasma, collected at different time points during the clinical management of GBM patients, and stored from two biobanks. Our longitudinal analysis unveils that vesiclemia is a dynamic parameter, which could inform on tumor burden and/or response to treatments. Further proteomic analysis unmasks the von Willebrand Factor (VWF) as a selective protein hallmark for GBM-patient separated EVs.

## Results

### Separation of Extracellular Vesicles from peripheral blood

Standardized, validated, and complementary procedures were implemented (14, 15) in order to isolate extracellular vesicles from plasma. As shown in Figure 1A, EV-contained fractions were collected through a low recovery/high specificity procedure using size exclusion chromatography (SEC), from a volume of 500 μL of 10,000g centrifugated plasma using an automatic fraction collector. Separated EV fractions were recovered by ultracentrifugation at 100,000g for two hours, resuspended in 0.22 μm-filtered PBS. To first evaluate the EV preparation degree of purity, the nature and extent of the protein cargo were estimated (Figure 1B). While an ascending concentration of proteins, most likely from bloodstream origin, was measured in the discarded fractions (qEV9-12), protein concentration was below the detectable levels in qEV7 and qEV8 fractions, discarding a massive extravesicular protein contamination in the eluted EVs (Figure 1B). Next, western-blot analysis shows the prominent presence of the EV-associated transmembrane tetraspanin CD9 in EV fractions, which was lost in later fractions (Figure 1C). Conversely, the Golgi protein marker GM130 was absent, excluding potential co-isolation of cellular components and debris (Figure 1C). We further documented the quality of separated EVs with cryo-electron microscopy (cryo-EM, Figure 1D). Large fields did not unmask massive aggregate contamination. Instead, typical lipid bilayer vesicles with a mean diameter of around 130 nm were detected. These electron-dense materials identified small EVs with circularity index of around 0.91, suggesting their integrity (Figure 1D). Likewise, plasmatic EVs were visualized with single particle tracking using interferometry light microscopy (ILM) technology, (Videodrop, Myriade), which enables the estimation of the concentration of particles according to Brownian motion (Figure 1E). The amount of EVs was readily detected in the qEV7 and qEV8 fractions with approximately 70 and 80 particles tracked per frame (orange circles), respectively, and a mean size of around 100-110 nm, as compared to filtered PBS negative control (Figures 1E-F). Likewise, the presence of EVs in both qEV7 and qEV8 fractions was underlined through ELISA detection of the tetraspanin transmembrane EV passenger CD63 (Figure 1H). We found an elevated level of CD63-positive particles, 5.5×10^9^ and 6.3×10^9^ particles/mL, in qEV7 and qEV8 fractions, respectively, as compared to 0.22 μm-filtered PBS used as a negative control (Figure 1H). These results were corroborated using flow cytometry detection of CD63-positive particles separated with anti-CD9 capture-beads (Figure 1I). Together, these findings confirm that plasmatic EVs could be separated and characterized with standard protocols, thereby ensuring the reliability of measuring vesiclemia.

**Figure 1.**
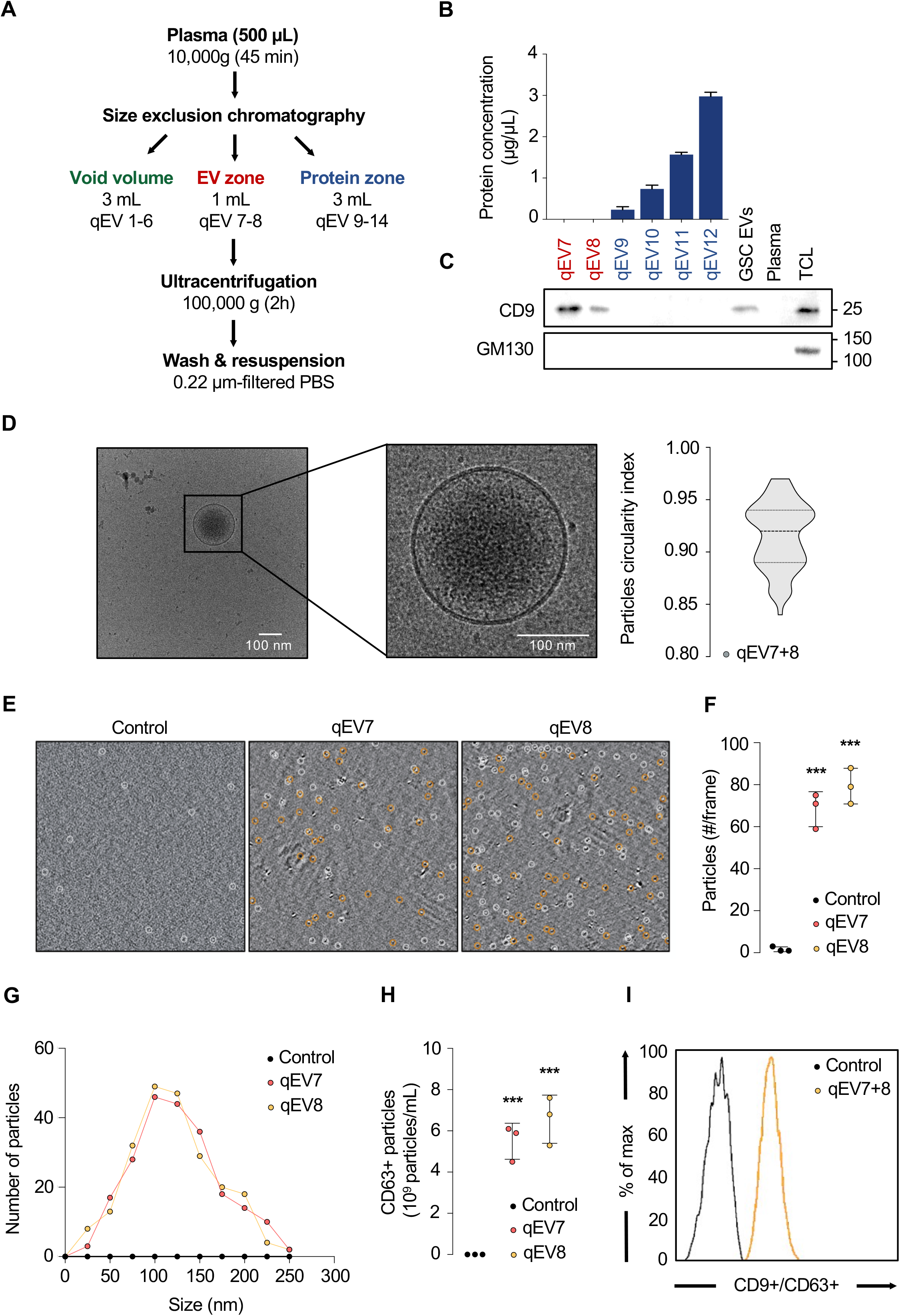
Separation of extracellular vesicles from human peripheral blood. **(A)** Diagram representing the separation of extracellular vesicles (EVs) from peripheral blood. **(B)** Total protein concentrations were measured in each size-exclusion chromatography (SEC) lysed fractions from qEV7 to qEV12. **(C)** Equal volume of protein lysates of qEV fractions were separated by SDS-PAGE and analyzed by immunoblot for CD9 (specific EV marker) and GM130 (putative intracellular membrane protein contaminant). EVs from GSC-conditioned media and total cell lysate (TCL) serve as controls. **(D)** Cryo-transmission electron microscopy (cryo-TEM) was deployed to image EVs in pooled qEV7 and qEV8 fractions and estimate sample purity (left panel). Nanoparticle morphology was evaluated using circularity index in n>70 cryo-TEM images (right panel). **(E-F)** Quantitative analysis of EVs was performed using interferometry light microscope (ILM), which tracks particles based on Brownian motion, in qEV7 and qEV8 fractions. Similarly, diameter distribution was monitored in representative qEV7 and qEV8 fractions. **(G)** EV abundance was estimated via CD63 ELISA. **(H)** Expression of CD63 was estimated by flow cytometry, following CD9-positive immunocapture beads in the fractions of interest. Data are representative of at least 3 independent experiments. ANOVA, ***p<0.001.

### Vesiclemia evolves along glioblastoma progression

To tackle the question whether quantity and/or quality of circulating EVs could represent potential hallmark of glioblastoma (GBM) evolution, we performed a longitudinal analysis of the vesiclemia (*i.e*. concentration of particles/mL) throughout tumor management. In this purpose, two bio-collections of plasmatic samples from GBM patients were harnessed, namely the French Glioblastoma Biobank (FGB), composed of multicentric samples collected upon diagnosis, and the Integrated Center for Oncology longitudinal biocollection (ICO) with peripheral blood sequentially collected alongside with tumor management (*i.e*. throughout the Stupp *et al*. protocol, during the follow-up and at tumor relapse) (Figure 2A). Table 1 reported the clinical information on the 30 patients enrolled here from two biobanks, including 20 at diagnosis, and 10 in longitudinal cohort, with 15- and 26-month median survival, respectively (Figure 2B). Plasmatic EVs were separated by size exclusion chromatography (SEC) and the vesiclemia was estimated either by single particle tracking using either ILM technology or ELISA for CD63.

**Figure 2.**
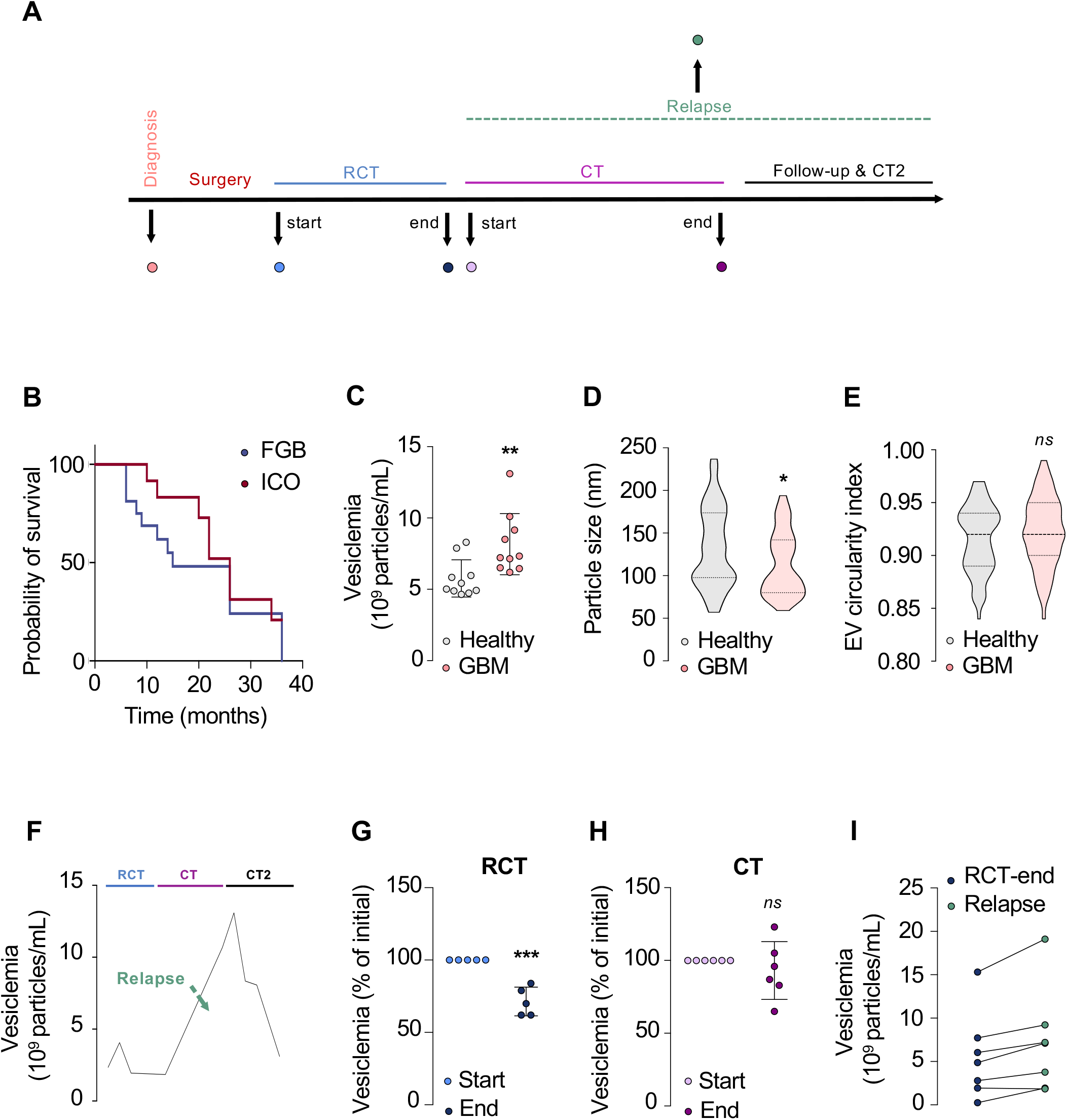
Vesiclemia evolves along glioblastoma progression. **(A)** Diagram of glioblastoma (GBM) management according to Stupp *et al*. protocol. RCT: radiochemotherapy, CT: chemotherapy. **(B)** Kaplan-Meier survival curves of both exploited biocollections from the French Glioblastoma Biobank (FGB) and Integrated Center for Oncology (ICO), with 20 and 10 patients, respectively. **(C)** Vesiclemia (number of particles/mL) was measured by interferometry light microscope (ILM) in healthy donors and GBM patients (n=10 in each group). **(D-E)** Analysis of plasmatic EV mean size in healthy donors and GBM patients using cryo-electron microscopy (cryo-TEM), n>65 for each group. Alternatively, nanoparticle morphology was evaluated using circularity index in cryo-TEM images. **(F-I)** Vesiclemia was measured via CD63 ELISA on the plasmatic fractions. Longitudinal vesiclemia in one patient throughout tumor management from Stupp protocol (RCT and CT) to second line CT2 (Bevacizumab) (F). Vesiclemia was measured in longitudinal samples, in order to assess the impact of radio-chemotherapy (RCT) (n=5) (G), chemotherapy (n=6) (H), and relapse (n=7) (I). Mann-Whitney test, *p<0.05, **p<0.01, ***p<0.001.

**Table 1.** Clinical information from GBM patients regarding age at diagnosis, gender, time to death and therapeutical care.

| Biobank | Age | Gender | Elapsed Time to Death | Surgery | RCT | 2<sup>nd</sup> Line |
| --- | --- | --- | --- | --- | --- | --- |
| FGB#1 | 61 | Female | 12 | partial | yes | n/a |
| FGB#2 | 71 | Male | 6 | total | yes | n/a |
| FGB#3 | 64 | Female | 8 | biopsy | yes | n/a |
| FGB#4 | 69 | Male | 9 | total | yes | Bevacizumab |
| FGB#5 | 74 | Male | 13 | total | yes | Bevacizumab |
| FGB#6 | 69 | Male | 6 | biopsy | yes | n/a |
| FGB#7 | 60 | Female | 15 | total | yes | Lomustine |
| FGB#8 | 72 | Male | 6 | total | yes | Irinotecan |
| FGB#9 | 64 | Male | 38 | total | yes | Irinotecan |
| FGB#10 | 69 | Female | 26 | total | yes | Bevacizumab |
| FGB#11 | 46 | Male | 12 | total | yes | Bevacizumab |
| FGB#12 | 55 | Male | 8 | total | yes | n/a |
| FGB#13 | 81 | Male | 13 | total | yes | n/a |
| FGB#14 | 63 | Female | 15 | biopsy | yes | Lomustine |
| FGB#15 | 66 | Male | 7 | biopsy | yes | n/a |
| FGB#16 | 65 | Male | 15 | total | yes | n/a |
| FGB#17 | 51 | Male | 17 | total | yes | Bevacizumab |
| FGB#18 | 74 | Female | 3 | partial | yes | n/a |
| FGB#19 | 48 | Male | 3 | total | yes | Bevacizumab |
| FGB#20 | 68 | Male | 14 | partial | yes | n/a |
| ICO#1 | 52 | Female | 26 | total | yes | Temozolomide |
| ICO#2 | 51 | Female | 24 | total | yes | Lomustine |
| ICO#3 | 67 | Male | 18 | total | yes | Bevacizumab |
| ICO#4 | 79 | Male | 30 | biopsy | yes | Bevacizumab |
| ICO#5 | 67 | Female | 20 | partial | yes | Bevacizumab |
| ICO#6 | 63 | Male | Alive | total | yes | n/a |
| ICO#7 | 77 | Female | 4 | biopsy | yes | Bevacizumab |
| ICO#8 | 55 | Male | Alive | partial | yes | Bevacizumab |
| ICO#9 | 64 | Male | 20 | biopsy | yes | Lomustine |
| ICO#10 | 79 | Male | 8 | partial | yes | n/a |

Two cohorts were used here (FGB and ICO, as described in method section). Elapsed time from diagnosis to death is expressed in months.

We firstly compared the vesiclemia of 10 newly diagnosed GBM patients and 10 healthy donors using interferometry microscopy. A significantly higher vesiclemia was found in the tumor patient group with a mean concentration about 8.5±×10^9^ particles/mL, as compared to healthy donors with 5.9±×10^9^ particles/mL, corroborating earlier findings (4) (Figure 2C). Moreover, electron microscopy analysis demonstrated plasmatic EVs from GBM patients upon diagnosis to be significantly smallest with a mean size of 101 nm, in comparison to the non-tumor donor group exhibiting a mean size of 129 nm (Figure 2D), while sharing similar circular morphology (Figure 2E).

To illustrate this longitudinal analysis, vesiclemia was measured in ten GBM patients and is presented here for one patient with conventional clinical features (1). in terms of administered treatments and time to relapse and demise (Table 1, Figure 2F). This personalized analysis highlights that this selected patient’s vesiclemia varies upon disease progression, with a noticeable peak at the relapse period. To further investigate how treatments might impact this biological parameter, we then explored the effect of the initial radiochemotherapy (RCT) on the plasmatic EV abundance of 5 patients sampled prior to and during RCT. Remarkably, all 5 patients displayed a drop of the vesiclemia during the treatment with a significant decrease of about 35%, outlining the impact of the RCT on the quantity of circulating plasmatic EVs (Figure 2G). Furthermore, we monitored the evolution of the vesiclemia during the 6-month temozolomide chemotherapy (CT), typically administered after one month break following RCT. The plasmatic EV abundance of 6 patients at the beginning (months 1-2) and the end (months 5-6) of the chemotherapy was not obviously modified over the treatment period (Figure 2H). Finally, the chronological evolution of the plasmatic EV abundance throughout the Stupp *et al*. protocol was evaluated for 7 relapsing patients, *i.e*. comparing values obtained at the initial RCT and at the time the tumor relapsed (Table 1). Noticeably, 6 out of these 7 patients unveiled an increase of about 40% of their vesiclemia when the tumor recurs, outlining a potential correlation between the plasmatic EV concentration and GBM relapse (Figure 2I). Therefore, these findings highlight the vesiclemia as a variable and dynamic biological parameter that might be regulated by several therapeutic factors, including radiations and drug administration, while placing the longitudinal analysis of the vesiclemia as a potential aid to anticipate patient follow-up and/or confirm tumor relapse. However, larger cohorts and longitudinal analyses of the evolution of the circulating EV cargo throughout tumor management are required to underpin the promising role of plasmatic EVs as biomarkers for glioblastoma progression.

### Proteomic analysis of circulating EVs from GBM patients unveils specific cargos

In order to get qualitatively insights on circulating EVs, label-free liquid chromatography tandem mass spectrometry was performed on EV fractions from 6 random plasmas from GBM patients (n=3) and healthy donors (n=3) (Figure 3A, Supplemental Table S1). Their cargo reached 80% and 74% of enriched EV proteins, when compared to established Exocarta and Vesiclepedia databases, respectively (Figure 3B). Further exploration of the database for annotation, visualization and integrated discovery (DAVID) highlighted “exosome” as the top Gene Ontology (GO) functions in plasma EV proteome, together with other dynamic processes, such as “cytoskeleton”, “adhesion” and “endoplasmatic reticulum functions” (Figure 3C). Principal component analysis (PCA) allowed to partition two distinct populations, grouping EV-protein cargo from GBM patients on one side and EV-protein cargo from healthy donors on the other (Figure 3D). In total, 291 and 189 proteins were detected within the three samples of each group, GBM patients and donors, respectively. A large part of them were shared between groups (*i.e*. 177 proteins, Figure 3D). Of note, a very few were exclusive to each group, namely 1 in the donor group (ANXA7) and 7 in the GBM group (PDLIM1, ANK1, EPB42, CALD1, CTTN, TMEM40, HSD17B10) (Figures 3D-E, Supplemental Table S2). Likewise, differentially expressed proteins were visualized on Volcano’ plot (Figure 3F, Supplemental Table S3). Heatmap unmasked 6 proteins significantly under-represented (TAGLN2, CSTA, YWHAG, YWHAB, YWHAE, and PPIA) in EVs from GBM patients, as compared to the ones from healthy donors (Figure 3G). Conversely, 2 proteins, namely VWF and FCN3, were detected as more abundant in EVs from GBM patients (Figure 3G). Data mining of The Cancer Genome Atlas (TCGA) further unveiled that the RNA expression of *VWF*, but not of *FCN3*, was heightened in GBM patients (Figure 3H). Moreover, several mutations were reported in the *VWF* gene in GBM patients, as well amplification, pointing again to the clinical interest of monitoring VWF expression (Figure 3I). In this context, VWF protein was significantly higher in EVs from GBM patient blood (mean concentration 59.6 ng/mL, n=20) than from healthy donors (mean concentration 45.7 ng/mL, n=20), at the time of diagnosis (Figure 3J). Our data thus support the notion that EVs from GBM patients are enriched with selective protein cargos that can be further surveyed in circulating EVs, together with vesiclemia.

**Figure 3.**
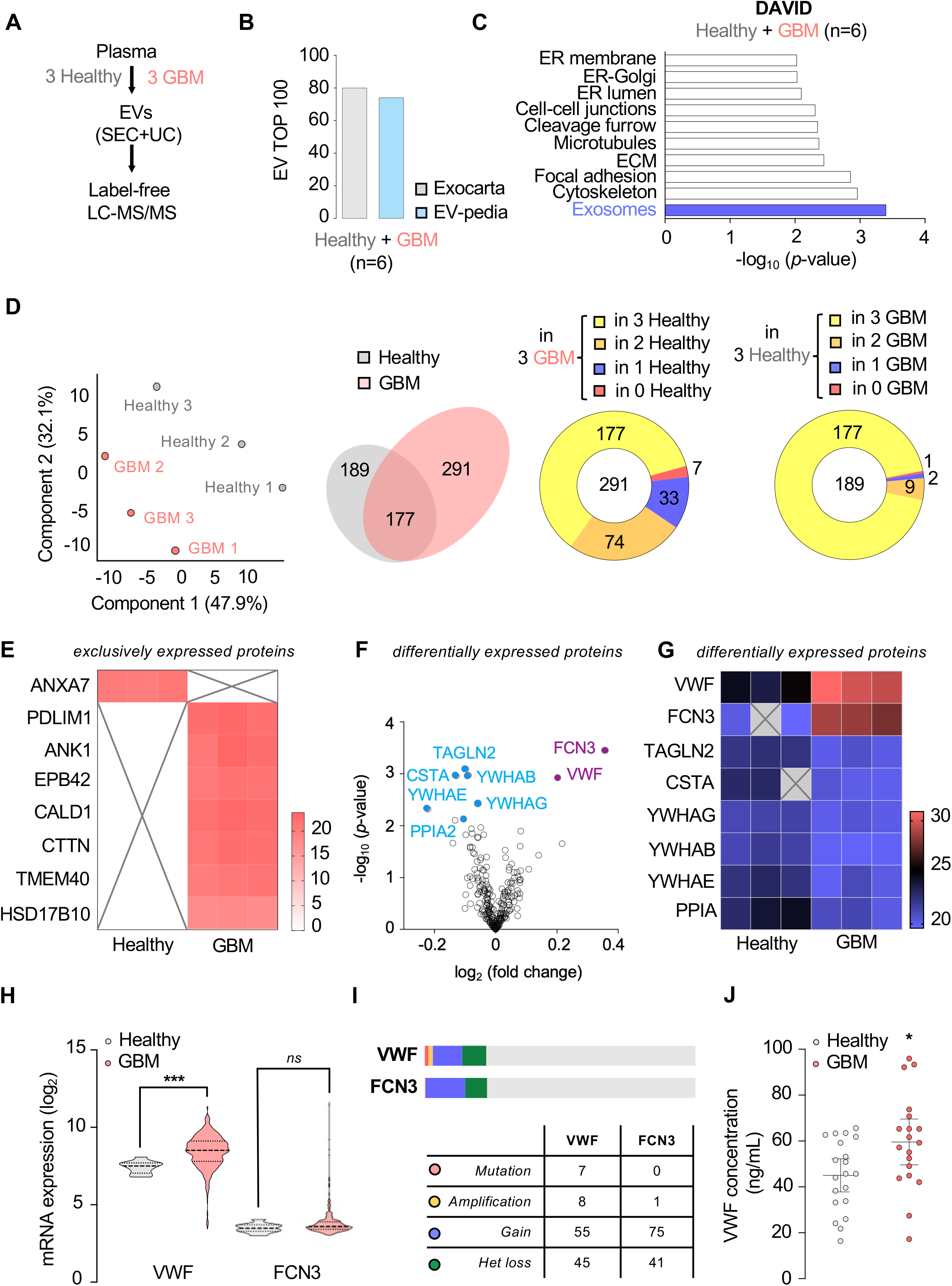
Proteomic analysis of circulating EVs from GBM patients unveils specific cargos. **(A)** Plasmatic extracellular vesicles (EVs) were separated from six plasmas (3 healthy donors and 3 GBM patients), as described in Figure 1A. Protein cargoes were quantitatively analyzed through label-free liquid chromatography tandem mass spectrometry (LC-MS/MS). **(B)** Representation of the top 100 EV-enriched proteins estimated with Exocarta and EVpedia databases, when combining proteomic analysis from 6 samples. **(C)** DAVID analysis of the top Gene Ontology (GO) functions from the 6 analyzed samples. **(D)** Principal component analysis of healthy donors and GBM patient samples (left panel). Venn diagram of the detected proteins and their repartition between healthy donor and GBM patient groups (middle and right panel). **(E)** Heatmap analysis of exclusively expressed proteins. **(F, G)** Volcano plot and heatmap analysis of differentially expressed proteins. **(H)** mRNA expression of von Willebrand factor (VWF) and ficolin-3 (FCN3) in healthy subjects and GBM patients estimated from The Cancer Genome Atlas (TCGA, n>500 for GBM patients). **(I)** Anomalies currently reported in VWF and FCN3 genes. **(J)** Concentration of VWF in plasmatic EVs from healthy donors and GBM patients (n=20). Mann-Whitney test, *p<0.05, ***p<0.001.

## Discussion

Seminal studies highlighted extracellular vesicles (EVs) as linchpin tools for tumor growth, survival and therapeutic refractoriness, as they carry and presumably disperse oncogenic material within the GBM ecosystem, as well as throughout the organism. Additionally, recent reports unveiled the presence of EVs into accessible body fluids including cerebrospinal fluid, plasma, and urine. In this context, circulating EVs have been thought to reflect valuable information about the evolving tumor while potentially act as a wide platform for non-invasive biomarkers that might assist in both GBM diagnosis and management. Notwithstanding, the evolution of plasmatic EV profile alongside tumor progression remains uncertain, making standardized information gathered at critical points of GBM evolution required.

We attempted here to assess circulating EVs as potential biomarkers of GBM. Often diagnosed at an advanced stage, there is not, nowadays, any biomarker able to screen for this pathology, to follow its evolution and relapse, and to monitor therapeutic efficacy and individual response to treatment. Recent works have highlighted potential translational applications of EVs in GBM. For instance, the specific RNA miR-21 was found in higher concentration in purified EVs from plasma and cerebrospinal fluid of patients with GBM (5, 16). Earlier studies proposed to block EV secretion and therefore halt tumor growth (17–20). The potential involvement of EVs in other cancers is also largely explored and provides a promising area of research. Notably, EVs has been suggested as markers in breast, pancreatic and prostate cancers (10). In this study, we brought a further comprehensive view on the quantitative and qualitative aspects of circulating EVs at different time course of the tumor management *(i.e*., upon diagnosis, during treatment and at relapse). Vesiclemia was reported in particles/mL and characterized by at least two different and complementary methods, including single particle analysis techniques (herein interferometry light microscopy) and biochemical techniques, such as ELISA. Electron microscopy, with close-up and wide field provided, was performed to complete EV characterization. According to a standardized and reproducible protocol for EV separation from peripheral blood, we firstly reiterated that the vesiclemia was higher in newly diagnosed GBM patients in comparison with healthy subjects (3, 4). Moreover, plasmatic EVs were found to be significantly smaller in tumor patients, while sharing similar circular morphology. We further highlighted a quantitative impact of the initial radio-chemotherapy on circulating EVs associated with a stabilizing effect of the six-month chemotherapy on the vesiclemia. Strikingly, our results stressed out a potential correlation between plasmatic EV abundance and GBM relapse, as 6 out of 7 patients displayed a rise of the vesiclemia upon tumor recurrence. Thus, our data support the idea that circulating EV profile quantitatively fluctuates throughout tumor natural evolution, corroborating recent works from Osti *et al*. where vesiclemia was found to be reduced after resective surgery while increased upon tumor relapse (4). Combined, our data strengthen the discovery that circulating EV load might reflect tumor burden. However, larger cohorts are essential to underpin these promising findings. Furthermore, standardized protocols of sampling, storage and analysis must be developed to ensure the reproducibility of the experiments and achieve robustness compatible with clinical routine.

In this work, proteomic analysis unmasked specific cargos in circulating EVs from GBM patients. Indeed, while principal component analysis demonstrated a clear segregation between tumor and healthy samples, a majority of proteins was shared by both conditions with two of them, namely von Willebrand factor (VWF) and ficolin-3 (FCN3), found to be significantly higher in GBM patients. However, only VWF mRNA expression was found elevated in GBM patients when interrogating TCGA databases, in contrast to FCN3 expression. This is in agreement with earlier findings that also highlight the theranostic value of VWF in brain tumor patients (21, 22). In line with these findings, VWF was significantly enriched in plasmatic EVs from GBM patients in comparison to healthy subjects. This could reflect not only EV-mediated tumor pro-angiogenic signaling (6, 7), but also potentially underlie the clinical pro-thrombotic status of cancer patients. To further assess circulating EVs as a non-invasive biomarker of GBM progression, proteomic analysis must be completed with transcriptomic, metabolomic and lipidomic studies to gain more insights on plasmatic EV qualitative aspect and its variation alongside tumor management. Moreover, the identification of a specific molecular signature discriminating plasmatic EVs directly emanating from GBM cells and the EV fractions resulting from pathophysiological release would represent a major step forward. However, this purpose remains complex and highly challenging, as for now no validated marker can track tumor EVs when unleashed in the bloodstream. Finally, functional studies need to address the role of circulating EVs on different cell types composing GBM microenvironment and brain parenchyma, such as endothelial cells, neurons and astrocytes. This should help in establishing EVs as key messengers hijacked by both tumor and stromal cells to the tumor own benefit.

In conclusion, our findings bring new insights on the characterization of circulating EVs from GBM patients and highlight plasmatic EVs as highly promising candidates for GBM bench-to-bedside research.

## Methods

### Ethic statement and plasma samples

Informed consents were obtained from all patients prior to sample collection in the frame of their medical exams. This study was reviewed and approved by the institutional review boards of Angers Hospital (CHU Angers, France) and Integrated Center for Oncology (Saint Herblain, France) and performed in accordance with the Helsinki protocol. Two biocollections were established and registered with plasmatic samples at different time course of glioblastoma progression. For the French Glioblastoma Biobank (#1476342v2, CHU Angers, France), liquid biopsies from glioblastoma patients were sampled at the time of diagnosis (23). For the Integrated Center for Oncology cohort (DC-2015-2457, ICO, Saint Herblain, France), 10 patients with glioblastoma were sequentially sampled at critical points of the Stupp *et al*. protocol (*i.e*., after resective surgery and upon radio-chemotherapy), during the follow-up and at relapse (24). Plasma samples were aliquoted and stored at −80°C. In addition, plasma samples from healthy donors were provided by the Etablissement Français du Sang (PLER-NTS2018-021, EFS, Nantes, France). For all samples, blood was collected on EDTA tubes and centrifuged at 1000g for 15min prior plasma storage at −80°C.

### The Cancer Genome Atlas (TCGA) analysis

The Cancer Genome Atlas (TCGA) database was explored via the Gliovis platform (http://gliovis.bioinfo.cnio.es/) (25). We interrogated RNAseq data from glioma patients for *VWF and FCN3* expression. Mutations, amplification, gain and heterozygosity loss are also reported.

### Extracellular vesicle separation

Extracellular vesicles (EVs) were separated from centrifugated plasma (10,000g, 20 min, 4°C) by size exclusion chromatography (SEC) using 70 nm Original qEV columns associated with an automatic fraction collector (Izon Science), according to the manufacturer’s protocol and recommendations from the International Society for Extracellular Vesicles (ISEV) (15). Briefly, plasma was centrifuged at 10000g for 10min before loading SEC columns. Of note, qEV7 and qEV8 fractions of 500 μL each were collected, pooled and harvested for further analysis. Alternatively, fractions were concentrated by ultracentrifugation (cryo-EM and proteomics). Samples were ultracentrifugated at 100,000g for 2 hours using OPTIMA MAX XP ULTRACENTRIFUGE with MLA-130 fixed-angle rotor (Beckman-Coulter) and pellets were resuspended in 0.22 μm-filtered PBS. All relevant experimental parameters were submitted to the open-source EV-TRACK knowledgebase (EV-TRACK.org)(13). EV-TRACK ID is EV210089.

### Quantitative analysis of EVs

According to recommendations from the ISEV (15), number of EVs was estimated through two different and complementary procedures. The number of EVs was measured using single particle tracking (Interferometry Light Microscopy, Videodrop, Myriade). Second, the abundance in EVs was also quantified by a biochemical technique using Exo-ELISA CD63 kit (EXEL-ULTRA-CD63-1, SBI) according to the manufacturer’s protocol. Technical triplicates (ILM) and duplicates (ELISA) were performed.

### Cryo-transmission electron microscopy analysis

EVs were separated and concentrated from 1.5 mL of plasma as described above, and then analyzed by cryo-transmission electron microscopy (Microscopy Rennes Imaging Center, Université de Rennes 1, France) using 200 kV Tecnai G2 T20 Sphera microscope (Field Electron and Ion Company), equipped with USC 4000 camera and a single axis cryo-holder model 626 (Gatan Microscopy). Both EV size and circularity index were estimated using ImageJ software.

### Flow cytometry analysis

EVs were separated and concentrated from 1 mL of plasma as described above and then analyzed by flow cytometry using CD9 exosome capture beads (ab239685, Abcam) according to the manufacturer’s protocol. PE anti-human CD63 staining (10896786, BioLegend) was next performed. CD9-conjugated bead-coupled EVs were harvested by centrifugation and resuspended in 0.22 μm-filtered PBS. Analysis was performed in triplicate using BD FACS-CANTOII (CYTOCELL platform, SFR François Bonamy, France) with 10,000 events recorded for each preparation. Histograms were mounted with FlowJo software.

### Immunoblotting

EVs were separated and concentrated from 500 μL of plasma as described above and the pellet was directly lysed in boiling Laemmli for 10 minutes. Proteins were resolved by SDS-PAGE, transferred onto nitrocellulose membranes and blotted with the following antibodies: CD9 (System Bioscience) and GM130 (Abcam) diluted at 1/1000. Membranes were incubated with HRP-conjugated secondary antibody diluted at 1/5000 and then revealed by chemiluminescence. Acquisitions were performed with Fusion software (Vilbert Lourmat).

### Mass spectrometry and proteomic analysis

Label free mass spectrometry analysis was performed (Proteom’IC, Institut Cochin, 3P5, Université Paris Sorbonne, Paris, France). EVs were separated and further concentrated from 1 mL of randomly chosen plasma of 3 GBM patients and healthy donors, as described above. Proteins were lysed and denatured in SDS 2%, 50mM Tris pH=8 (5 min at 95 °C), hydrolyzed with trypsin and released peptides were analyzed in triplicate by liquid chromatography with tandem mass spectrometry (LC-MS/MS) using Orbitrap Fusion mass spectrometer (ThermoFisher Scientific). Obtained spectra were processed using MaxQuant and Perseus servers and compared to Swiss-Prot/TrEMBL banks for protein identification. Label-free quantification was performed on three samples in the control and GBM groups, while FDR and number of identified peptides are reported (Supplemental Table 1). Number of valid values (VV) per group (0, 1, 2 or 3) allows the selection of proteins that are considered as expressed (VV of 2 or 3) or absent (VV of 0 or 1). Principal component analysis (PCA) was next performed on 100% VV proteins, while peptides with a p-value inferior to 0.01 (t-test) were considered differentially expressed.

### Quantitation of von Willebrand factor (VWF)

VWF concentration was estimated on EV preparation using human VWF ELISA kit (EHVWFX, ThermoFisher Scientific) according to the manufacturer’s protocol. Technical replicates by serial dilution were performed.

### Quantitation of proteins

Protein concentration of each qEV fraction was estimated by BCA protein assay according to the manufacturer’s protocol (ThermoFisher Scientific). Technical replicates by serial dilution were performed.

### Statistical analysis

Statistical analyses were performed with Prism8 software using unpaired two-tailed Mann-Whitney test (non-parametric test) and ordinary one-way analysis of variance (ANOVA). For all the experiments, a p-value inferior to 0.05 in considered significant.

## Author contributions

conception and design (QS, JSF, JG), acquisition of data, analysis and interpretation of data (QS, GAG, CAN, NB, LG, JSF, JG), drafting the article (QS and JG). All authors approved the manuscript.

## Acknowledgments

We thank SOAP team members. We are also grateful to Cytocell core-facility (SFR Santé François Bonamy, Nantes, France) and to Aurelien Dupont for cryo-TEM analysis (MRIC, Rennes, Nantes). We also thank Francois Guillonneau, Cédric Broussard and Virginie Salnot from Plateforme Proteomique 3P5, Universite de Paris Sorbonne, Institut Cochin, Paris, France for performing sample preparation and data acquisition and analysis. We would like also to acknowledge the French Glioma Biobank, Angers CHU Angers, France) and CRB-Tumorotheque de l’ICO (ICO, Saint Herblain, France).

This research was funded by Fondation pour la Recherche Médicale (Equipe labellisée DEQ20180339184), Fondation ARC contre le Cancer (PJA20171206146 and PJA20191209477), Institut National du Cancer (2019-151, 2019-291), Ligue nationale contre le cancer comités de Loire-Atlantique, Maine et Loire, Vendée, Ille-et-Vilaine (JG). QS received Master fellowship from Plan Cancer 2014-2019. The team is part of the SIRIC ILIAD (INCA-DGOS-Inserm_12558).

## References

1. Stupp R et al. Radiotherapy plus concomitant and adjuvant temozolomide for glioblastoma. N. Engl. J. Med. 2005;352(10):987–996.

2. Sabbagh Q, Andre-Gregoire G, Guevel L, Gavard J. Vesiclemia: counting on extracellular vesicles for glioblastoma patients. Oncogene 2020;39(38):6043–6052.

3. Andre-Gregoire G, Bidere N, Gavard J. Temozolomide affects Extracellular Vesicles Released by Glioblastoma Cells. Biochimie 2018;155:11–15.

4. Osti D et al. Clinical Significance of Extracellular Vesicles in Plasma from Glioblastoma Patients. Clin. Cancer Res. 2019;25(1):266–276.

5. Skog J et al. Glioblastoma microvesicles transport RNA and proteins that promote tumour growth and provide diagnostic biomarkers. Nat. Cell Biol. 2008;10(12):1470–1476.

6. Treps L, Perret R, Edmond S, Ricard D, Gavard J. Glioblastoma stem-like cells secrete the pro-angiogenic VEGF-A factor in extracellular vesicles. J. Extracell. Vesicles 2017;6(1):1359479.

7. Treps L et al. Extracellular vesicle-transported Semaphorin3A promotes vascular permeability in glioblastoma. Oncogene 2016;35(20):2615–2623.

8. André-Grégoire G, Gavard J. Spitting out the demons: Extracellular vesicles in glioblastoma. Cell Adhes. Migr. 2017;11(2):164–172.

9. van Niel G, D’Angelo G, Raposo G. Shedding light on the cell biology of extracellular vesicles. Nat. Rev. Mol. Cell Biol. 2018;19(4):213–228.

10. Becker A et al. Extracellular Vesicles in Cancer: Cell-to-Cell Mediators of Metastasis. Cancer Cell 2016;30(6):836–848.

11. Figueroa JM et al. Detection of wild-type EGFR amplification and EGFRvIII mutation in CSF-derived extracellular vesicles of glioblastoma patients. Neuro-Oncol. 2017;19(11):1494–1502.

12. Hoshino A et al. Extracellular Vesicle and Particle Biomarkers Define Multiple Human Cancers. Cell 2020;182(4):1044–1061.

13. Eitan E et al. Age-Related Changes in Plasma Extracellular Vesicle Characteristics and Internalization by Leukocytes. Sci. Rep. 2017;7(1):1342.

14. EV-TRACK Consortium et al. EV-TRACK: transparent reporting and centralizing knowledge in extracellular vesicle research. Nat. Methods 2017;14(3):228–232.

15. Théry C et al. Minimal information for studies of extracellular vesicles 2018 (MISEV2018): a position statement of the International Society for Extracellular Vesicles and update of the MISEV2014 guidelines. J. Extracell. Vesicles 2018;7(1):1535750.

16. Akers JC, Gonda D, Kim R, Carter BS, Chen CC. Biogenesis of extracellular vesicles (EV): exosomes, microvesicles, retrovirus-like vesicles, and apoptotic bodies. J. Neurooncol. 2013;113(1):1–11.

17. Ostrowski M et al. Rab27a and Rab27b control different steps of the exosome secretion pathway. Nat. Cell Biol. 2010;12(1):19–30.

18. Bobrie A et al. Rab27a supports exosome-dependent and -independent mechanisms that modify the tumor microenvironment and can promote tumor progression. Cancer Res. 2012;72(19):4920–4930.

19. Bronisz A et al. Extracellular Vesicles Modulate the Glioblastoma Microenvironment via a Tumor Suppression Signaling Network Directed by miR-1. Cancer Res. 2014;74(3):738–750.

20. Atai NA et al. Heparin blocks transfer of extracellular vesicles between donor and recipient cells. J. Neurooncol. 2013;115(3):343–351.

21. Marfia G et al. Prognostic value of preoperative von Willebrand factor plasma levels in patients with Glioblastoma. Cancer Med. 2016;5(8):1783–1790.

22. Navone SE et al. Significance and Prognostic Value of The Coagulation Profile in Patients with Glioblastoma: Implications for Personalized Therapy. World Neurosurg. 2019;121:e621–e629.

23. Clavreul A et al. The French glioblastoma biobank (FGB): a national clinicobiological database. J. Transl. Med. 2019;17(1):133.

24. Heymann, D et al. Centre de Ressources Biologiques-Tumorothèque: Bioresources and associated clinical data dedicated to translational research in oncology at the Institut de Cancérologie de l’Ouest, France. Open J Bioresour. 2020;(7):5.

25. Bowman RL, Wang Q, Carro A, Verhaak RGW, Squatrito M. GlioVis data portal for visualization and analysis of brain tumor expression datasets. Neuro-Oncol. 2017;19(1):139–141.

26. EV-TRACK Consortium et al. EV-TRACK: transparent reporting and centralizing knowledge in extracellular vesicle research. Nat. Methods 2017;14(3):228–232.

