## Supplemental tables for "The von Willebrand Factor stamps Plasmatic Extracellular Vesicles from Glioblastoma Patients"

| **name** | **peptide count** | **score** | **coverage (%)** |
| --- | --- | --- | --- |
| **ANAX7** | 4;3;5;0;0;0 | 6.9 | 12.2 |
| **PDLIM1** | 0;1;1;8;12;9 | 83.3 | 55.5 |
| **ANK1** | 0;0;0;4;20;18 | 67.3 | 13.2 |
| **EPB42** | 0;0;1;2;9;8 | 28.7 | 17.0 |
| **CALD1** | 0;0;2;10;16;11 | 49.8 | 33.1 |
| **CTTN** | 1;0;0;3;12;11 | 74.4 | 35.5 |
| **TMEM40** | 0;0;0;2;5;4 | 18.3 | 26.2 |
| **HSD17B10** | 0;0;1;2;2;2 | 4.2 | 9.6 |

**Supplementary Table S2.** Exclusively Expressed Proteins from the label-free proteomic analysis in 3 control (1;2;3) and GBM (1;2;3) plasmatic EV fractions.

| **name** | **peptide count** | **score** | **coverage (%)** |
| --- | --- | --- | --- |
| **TAGLN2** | 4;6;6;3;6;4 | 97.4 | 44.5 |
| **CSTA** | 5;4;0;4;2;3 | 16.5 | 67.3 |
| **YWHAG** | 5;7;8;6;8;8 | 67.5 | 35.2 |
| **YWHAE** | 11;13;13;7;10;9 | 36.4 | 55.7 |
| **YWHAB** | 8;11;10;8;8;9 | 47.0 | 48.8 |
| **PPIA** | 7;9;7;6;6;3 | 38.9 | 53.3 |
| **FCN3** | 3;1;3;13;13;12 | 85.4 | 53.8 |
| **VWF** | 26;27;57;99;101;93 | 323 | 44.0 |

**Supplementary Table S3.** Differentially Expressed Proteins from the label-free proteomic analysis in 3 control (1;2;3) and GBM (4;5;6) plasmatic EV fractions.
